# A near complete haplotype-phased genome of the dikaryotic wheat stripe rust fungus *Puccinia striiformis* f. sp. *tritici* reveals high inter-haplotype diversity

**DOI:** 10.1101/192435

**Authors:** Benjamin Schwessinger, Jana Sperschneider, William S. Cuddy, Diana P. Garnica, Marisa E. Miller, Jennifer M. Taylor, Peter N. Dodds, Melania Figueroa, Park F. Robert, John Rathjen

## Abstract

A long-standing biological question is how evolution has shaped the genomic architecture of dikaryotic fungi. To answer this, high quality genomic resources that enable haplotype comparisons are essential. Short-read genome assemblies for dikaryotic fungi are highly fragmented and lack haplotype-specific information due to the high heterozygosity and repeat content of these genomes. Here we present a diploid-aware assembly of the wheat stripe rust fungus *Puccinia striiformis* f. sp. *tritici* based on long-reads using the FALCON-Unzip assembler. RNA-seq datasets were used to infer high quality gene models and identify virulence genes involved in plant infection referred to as effectors. This represents the most complete *Puccinia striiformis* f. sp. *tritici* genome assembly to date (83 Mb, 156 contigs, N50 1.5 Mb) and provides phased haplotype information for over 92% of the genome. Comparisons of the phase blocks revealed high inter-haplotype diversity of over 6%. More than 25% of all genes lack a clear allelic counterpart. When investigating genome features that potentially promote the rapid evolution of virulence, we found that candidate effector genes are spatially associated with conserved genes commonly found in basidiomycetes. Yet candidate effectors that lack an allelic counterpart are more distant from conserved genes than allelic candidate effectors, and are less likely to be evolutionarily conserved within the *P. striiformis* species complex and *Pucciniales*. In summary, this haplotype-phased assembly enabled us to discover novel genome features of a dikaryotic plant pathogenic fungus previously hidden in collapsed and fragmented genome assemblies.

**Importance:** Current representations of eukaryotic microbial genomes are haploid, hiding the genomic diversity intrinsic to diploid and polyploid life forms. This hidden diversity contributes to the organism’s evolutionary potential and ability to adapt to stress conditions. Yet it is challenging to provide haplotype-specific information at a whole-genome level. Here, we take advantage of long-read DNA sequencing technology and a tailored-assembly algorithm to disentangle the two haploid genomes of a dikaryotic pathogenic wheat rust fungus. The two genomes display high levels of nucleotide and structural variations, which leads to allelic variation and the presence of genes lacking allelic counterparts. Non-allelic candidate effector genes, which likely encode important pathogenicity factors, display distinct genome localization patterns and are less likely to be evolutionary conserved than those which are present as allelic pairs. This genomic diversity may promote rapid host adaptation and/or be related to the age of the sequenced isolate since last meiosis.

## Introduction

The Basidiomycota and the Ascomycota constitute the two largest fungal phyla and contain many of the most damaging crop pathogens (1). The dominant life phase for most basidiomycete species is dikaryotic, where two haploid nuclei coexist within one cell (2). To date, about 475 basidiomycete fungal genome sequences representing some 245 species are available in the public domain (September 2017, https://www.ncbi.nlm.nih.gov/genome/). These genome references are either representations of the haploid life stage of a species (3), or collapsed and mosaic assemblies of the dikaryotic state (4–7). Hence, the information about the inter-haplotype variation in dikaryotic basidiomycota beyond single nucleotide polymorphisms (SNPs) and small insertions and deletions (INDELs) is very limited. The absence of haplotype-phased information limits the studies of genome architecture and evolution, particularly for the rust fungi of the order *Pucciniales*, many of which are extremely destructive pathogens of economically important crops including cereals, coffee, and soybean (8–13).

Stripe, stem and leaf rusts are the three rust diseases that impact wheat production, one of the most important staples in human diets (14). Of these, stripe rust caused by *Puccinia striiformis* f. sp. *tritici* (*Pst*) is currently the most damaging disease with estimated annual losses of $USD 1 billion (15, 16). As a biotrophic pathogen, *Pst* colonizes living hosts and extracts large amounts of nutrients from plant cells through specialized structures called haustoria. The large tax on host energy reserves caused by *Pst* infection results in yield losses mostly associated with poor grain filling (17).

The full life cycle of *Pst* involves asexual and sexual reproductive phases associated with the production of specific spore types (13, 17). The damage to wheat occurs during the asexual cycle and results from the repeated infections throughout the growing season, which cause exponential amplification of dikaryotic urediniospores. *Pst* infects more than 30 varieties of *Berberis* spp. and *Mahonia* spp. to complete its full sexual life-cycle that involves four additional spore stages and sexual recombination during meiosis. Sexual reproduction is restricted geographically to the Himalayan region (Nepal, Pakistan and China), where it leads to high levels of genetic diversity that are largely absent from other parts of the world. This makes the extended Himalaya region the center of *Pst* diversity and the main source for new highly virulent *Pst* isolates (12, 21).

Genetic resistance in the host plant, particularly race-specific resistance, is often used in the field to reduce damage by pathogenic rust fungi (22, 23). Race-specific resistance is generally conferred by dominant resistance (R) genes in the host which recognize specific avirulence (*Avr*) alleles within the pathogen. Mechanistically, *Avr* alleles encode variants of virulence effector proteins, and the *R* gene typically encodes a nucleotide-binding leucine rich repeat (NB-LRR) protein that detects the Avr protein within the infected plant cell. In case of *Pst,* more than 75 yellow rust resistance genes (*Yr*) have been catalogued to date. A given *Pst* isolate has a characteristic spectrum of *Avr* alleles that can be distinguished on a set of wheat tester lines containing these *Yr* genes (24). The collective virulence phenotypes on such differential set defines the *Pst* pathotype. Wheat stripe rust epidemics are associated with the appearance of genetically novel pathotypes which are not recognized by currently employed *R* genes and hence grow on commercial wheat cultivars. As such, incursions of exotic stripe rust isolates with new virulences can play a role in disease outbreaks, for instance the Warrior *Pst* lineage which invaded Europe in 2011 was highly successful as it was virulent on the wheat cultivars grown at that time (25, 26). In addition to this novel exotic incursion, it is well documented that *Pst* rapidly evolved new virulence traits on a continental scale in Australia following its introduction in 1979 (27). However, the mechanisms underlying the evolution of these new pathotypes remains understudied as no genetic locus contributing to the evolution of virulence has yet been identified in *Pst*. While new combinations of alleles generated during sexual recombination can lead to the emergence of new pathotypes, the contribution of other genetic and molecular events to pathogen evolution during asexual reproduction is unclear. Presumably, the occurrence of mutation explains the loss of *Avr* specificities and the adaptation to otherwise resistant wheat cultivars (13, 27).

Most agriculturally important fungi are haploid with small genomes (28). Rusts on the other hand are dikaryotic in the asexual phase and have expanded genomes with large amounts of repetitive sequence (6, 7). It is likely that the separation of rust genomes into two haploid copies contributes to their rapid evolution. Existing *Pst* genome sequences suffer from the use of short-read sequencing technologies, which prevents characterization of individual haploid genomes, while the high percentage of repetitive DNA reduces the size of contigs that can be assembled (4, 5, 29). The overall similar gene content of each genome causes the reads from allelic variants to collapse upon assembly, producing a consensus sequence that loses haplotype (phasing) information. Read mapping to the consensus reference reveals that the two genomes are highly heterozygous for SNPs (5, 7), but differences in effector and gene content are undetectable. These problems can be addressed to some extent by using traditional Sanger long-sequence reads or strategies such as fosmid-to-fosmid sequencing (6, 7), however, these approaches are expensive. Opportunities to resolve the questions at higher resolution have arisen from new technologies that generate very long sequencing reads (>10kb) (30, 31).

Here, we use long-read sequencing to provide a near-complete haplotype-phased genome assembly for an isolate representing the first pathotype of *Pst* detected in Australia in 1979 (27). Our assembly provides the most complete *Pst* genome reference to date with over 97% of all basidiomycete benchmarking universal single copy orthologs (BUSCOs) captured (32). In addition, phased haplotype information for over 92% of the genome enabled us to detect high inter-haplotype diversity at the nucleotide and structural level, which identifies allelic variation and that 25% of all genes lack a clear allelic counterpart. We identified over 1,700 candidate effector genes, which are more often spatially associated with each other and conserved BUSCOs rather than with repetitive elements. Non-allelic candidate effectors that lack counterparts in the alternate haploid genome region are less likely to be evolutionarily conserved in other rust fungi. Thus, the highly contiguous haplotype assembly has allowed discovery of novel genome features that may be linked to the rapid evolution of this devastating pathogen.

## Results and Discussion

### Haplotype-aware genome assembly of an Australian *Puccinia striiformis* f. sp. *tritici* isolate

The main aim of this study was to generate a high quality reference genome for *Pst*. For this purpose we sequenced a single pustule isolate of the Australian founder pathotype *Pst* 104E137A-, collected in 1982 (abbreviated as Pst-104E). We sequenced 13 PacBio SMRT cells obtaining a total of 13.7 Gb of data with an average read length of 10,710 bases, and a read length N50 of 15,196 bases (Table S1). We assembled these data using the diploid-aware assembler FALCON-Unzip (30) to obtain a synthetic haplotype-phased reference genome. The FALCON-Unzip assembler is designed to phase structural variations and associated single nucleotide polymorphisms (SNPs) into distinct haplotype blocks. This gives rise to a primary assembly (primary contigs) and linked haplotype blocks (haplotigs). The haplotigs represent the alternative genome structure with respect to primary contigs. FALCON-Unzip does not always link physically connected phase blocks, and primary contigs can represent sequences from either of the two haploid genomes (30).

Previous unphased *Pst* genome assemblies ranged in size between 53 and 115 Mb (4, 5, 7, 29). In an attempt to reconcile the differences in reported genome sizes, we used GenomeScope to estimate the haploid genome size using *k*-mer frequencies (30-mer) in two Illumina short-read datasets of Pst-104E (33). Based on this analysis we estimate a haploid genome size of 68-71 Mb, with a heterozygosity (SNPs and INDELs) rate of approximately 1.2%. We assembled our long-read data into 156 primary contigs with a total length of 83 Mb after manual curation. The corresponding phased haplotype blocks were contained in 475 haplotigs with a total size of 73 Mb (Table 1).

**Table 1:** Summary of *Pst*-104E genome assembly and annotation Summary statistics for the genome assembly according to the three different contig categories as described in the main text. ^a^Candidate effectors have been predicted based on the machine learning algorithm EffectorP and transcriptional upregulation during infection of wheat as described in the main text.

|  | Primary assembly |  | Haplotype assembly |
| --- | --- | --- | --- |
|  | Primary contigs with haplotigs | Primary contigs without haplotigs | Haplotigs |
| Number of contigs | 99 | 57 | 475 |
| Bases | 79,770,604 | 3,585,012 | 73,478,481 |
| TEs Coverage | 53.72 | 67.17 | 52.82 |
| Number of genes | 15303 | 625 | 14321 |
| Average gene length | 1210 | 1290 | 1189 |
| Average introns per gene | 3.45 | 2.70 | 3.42 |
| Number of genes per 10kb | 1.92 | 1.74 | 1.95 |
| BUSCOs | 1395 | 49 | 1293 |
| Number of BUSCOs per 10kb | 0.17 | 0.14 | 0.18 |
| Number of candidate effectors<sup>a</sup> | 1523 | 49 | 1390 |
| Number of candidate effectors<sup>a</sup><br>per 10kb | 0.19 | 0.14 | 0.19 |

These assembly statistics are a vast improvement over previous assemblies in terms of connectivity and number of contigs (Figure 1A). The primary assembly has a contig N50 of 1.3 Mb compared to a scaffold N50 of 0.5 Mb for *Pst*-78 or contig N50 of 5.1 kb for *Pst*-130, often referred to as the reference genome (4, 26, 29). In addition we identified 1,302 (97.5%) out of the 1,335 benchmarking genes (BUSCO v2; http://busco.ezlab.org/) (32) that are highly conserved in basidiomycetes, with only 10 (0.7%) missing in our combined assembly before filtering for genes related to transposable elements (TE). Our final assembly has 1,292 (96.8%) complete BUSCOs with 19 (1.4%) missing. This compares to wide variation in BUSCOs that can be identified from previous assemblies, ranging from 35.7% for *Pst-887* to 95.6% for *Pst-78* (Figure 1B). In summary, our assembly currently represents the most complete *Pst* reference in terms of contiguity, haplotype-phased information, and gene content. This advance provides a new resource to investigate genome architecture and inter-haplotype variation within this dikaryotic plant pathogen.

**Figure 1:**
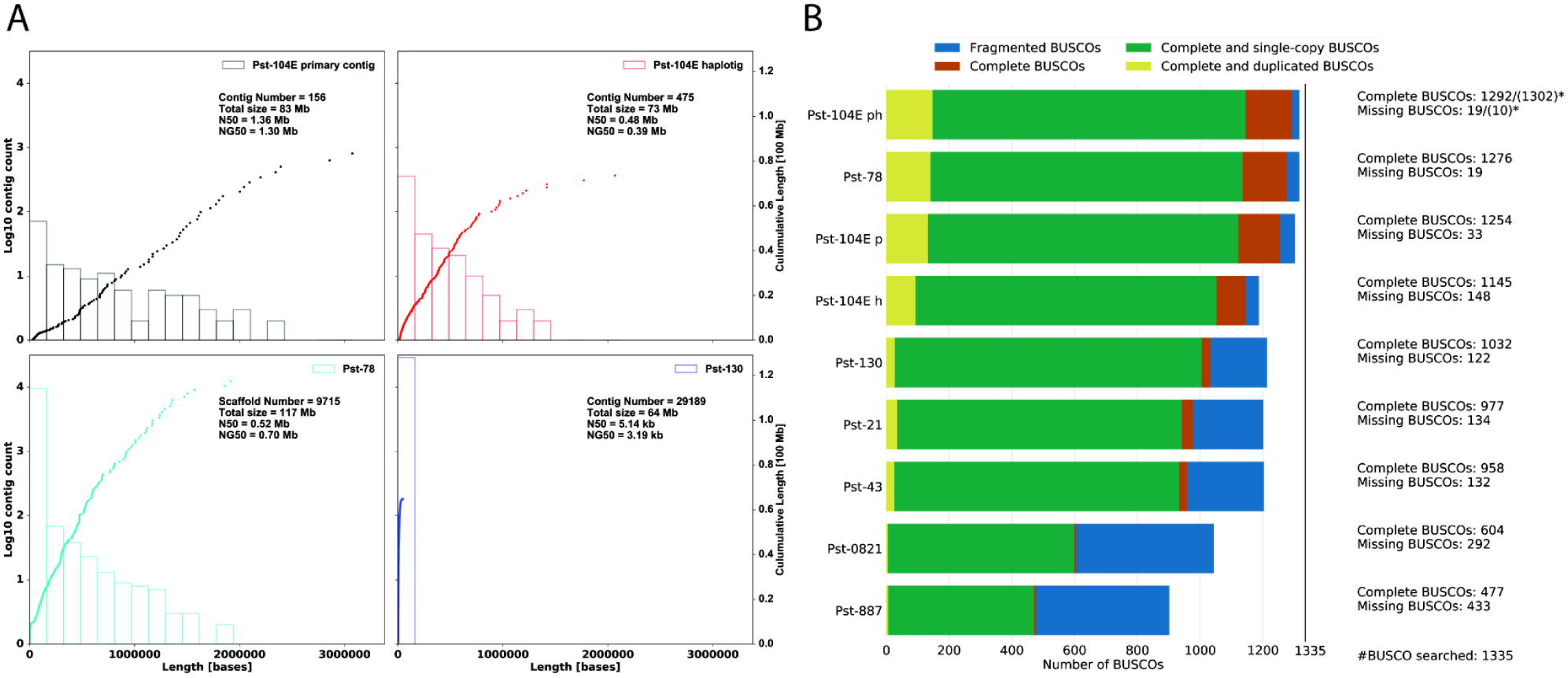
The *Pst*-104E genome assembly is highly contiguous and complete. A) Comparison of the Pst-104E primary and haplotig assemblies with the two most complete publicly available *Pst* genome assemblies, *Pst*-78 and Pst-130. The histograms and the left y-axis show log10 counts of contigs within each size bin. The dots and the right y-axis show the cumulative size of small to large sorted contig lengths. Each dot represents a single contig of given size shown on the x-axis. Each plot also shows the number of contigs or scaffolds, total assembly size, N50 of the assembly and NG50 assuming a genome size of 85 Mb. NG50 is the N50 of an assembly considering the estimated genome size instead of the actual assembly size. This enables comparisons between different sized assemblies. B) Genome completeness assessed using benchmarking universal single-copy orthologs (BUSCOs) for basidiomycota (odb9) as proxy. The graph shows BUSCO results for Pst-104E primary (p), haplotig (h) and non-redundantly combined (ph) assemblies in comparison to all publicly available *Pst* genome assemblies with gene models including *Pst*-78, *Pst*-130, *Pst*-21, *Pst*-43, *Pst*-0821, and *Pst*-887. The analysis was performed on the protein level using publicly available gene models. The * indicates the actual number of identified BUSCOs for the complete *Pst*-104E ph assembly before filtering gene models for similarity with genes encoded by transposable elements.

### High levels of inter-haplotype block variation

The *Pst*-104E primary assembly covers 83 Mb in a total of 156 primary contigs. Within this assembly, 99 primary contigs (~80 Mb) are associated with 475 haplotigs (~73 Mb), representing phased information for 92% of the primary contigs. These primary contigs are referred to as primary contigs with haplotigs. Overall, short-read mapping coverage analysis strongly supported our genome assembly. When we mapped short reads against the primary assembly, we observed a bimodal distribution of coverage. With a haploid genome coverage around ~60-fold and a diploid genome coverage at ~120-fold (Figure S1A). Regions with ~60-fold coverage are sequences that are distinct enough between the two haplotypes that only short reads originating from these specific sequences are able to map. Regions with ~120-fold coverage are sequences that are similar enough in the two haplotypes that short reads from both haplotypes collapse on the primary contig sequence when mapping against primary contigs only.

In contrast, when reads were mapped against both primary contigs and haplotigs, we found haplotigs and phased primary contig regions, that align to haplotigs, display ~60-fold coverage (Figure S1E and F). These are regions of the Pst-104E genome that are phased into two haplotype blocks. In addition, primary contig regions that lack an associated haplotig display mostly ~60-fold coverage (Figure S1C and G) suggesting that these are largely sequences specific to one haplotype and not collapsed highly similar regions of corresponding chromosome copies. Only a minor fraction of primary contigs show ~120-fold coverage (Figure S1D and G) when mapping against primary contigs and haplotigs, indicating the presence of a low residual of unphased sequences in our assembly.

Of the 57 primary contigs (~3.6 Mb) without associated haplotig (Table 1) 51 (~3.4 Mb) are likely single-haplotype-specific sequences because they display similar mean read coverage (~60-fold) to phased haploid regions of the genome (Figure S1). This high level of phasing enabled us to investigate inter-haplotype variation at a whole-genome scale. Previous studies using Illumina short-reads mapped against the consensus merged haplotype assemblies estimated *Pst* inter-haplotype variation based on heterozygous SNPs at between 0.5% - 1% (5, 7, 29). Taking a similar approach, we identified approximately 0.5% (416,460 heterozygous SNPs) of the genome as variable when mapping lllumina short reads against primary contigs only. However, we estimated a dramatically higher level of inter-haplotype variation when using this phased assembly. For this analysis, we aligned all haplotigs with their corresponding primary contigs and estimated variations using Assemblytics (34, 35). Assemblytics defines six major categories of structural variations including insertions and deletions, tandem repeats identified by overlapping alignments and other types of repeats suggested by gapped non-unique contig alignments (see Figure 2A for illustration of the six different variant categories) and divides these according to size in bins (Figure 2A). This analysis revealed that structural variation comprised 6.4% (~5.10/79.77 Mb) of the primary assembly space when compared to respective haplotigs (Figure 2A) (34). The variation between two primary contigs and their respective haplotigs is illustrated in the dot plots shown in Figure 2B and C, which visualizes large-scale inversions, deletions and insertions in haplotigs associated with two primary contigs. It is likely that the actual difference between the two haplotypes is higher than the estimated 6%, because calculations were restricted to a maximal variant size of 10 kb and do not include primary contigs without haplotigs, which account for another ~3.6%. Overall, the dramatic difference in estimated inter-haplotype variation between previous assemblies (5, 7, 29) and short-read based prediction programs (33) is likely caused by the fact that most of the observed variations are contained in size bins greater than 500 bases, which is not detectable with Illumina short-read data and highly fragmented assemblies.

**Figure 2:**
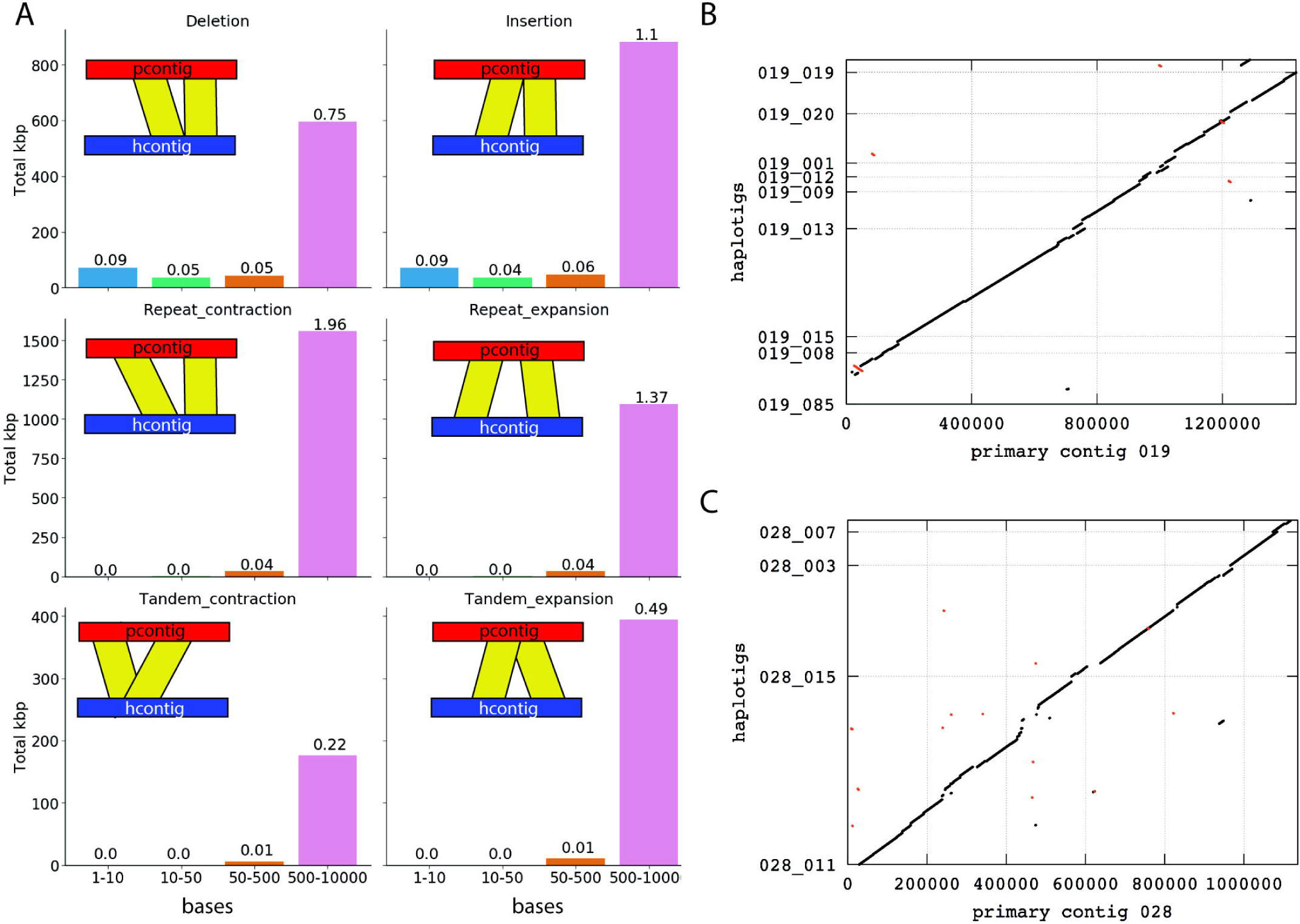
The *Pst-104E* genome is characterized by high levels of inter-haplotype variation. A) Summary of inter-haplotype variation between primary contigs and their respective haplotigs using Assemblytics. Each plot indicates the number of bases that are spanned by the specific variation category, which is illustrated by a cartoon. The number labelling each histogram represents the % of the total size of primary contigs with haplotigs that are contained within this variation type and size bin. B) and C) Two representative whole genome alignments of primary contigs 019 and 028 with their respective haplotigs. This illustrates the large-scale variations summarized in A).

### Over half of the *Pst-104E* genome is covered by repetitive sequences

We annotated primary contigs and haplotigs independently based on our observations of high levels of heterozygosity between the two (Figure 2 and S2). We first identified and classified transposable elements (TEs) using the REPET pipeline (36) to the order level based on the Wicker classification (37). We further transferred superfamily annotation from the underlying BLAST (38) hits if they agreed with the REPET annotations and with each other. There was no major difference between TE coverage of primary contigs 54% (~45 Mb) and haplotigs with 53% (~39 Mb) (Figure S2). However, primary contigs that lacked haplotigs had a larger proportion of TEs with a total coverage of 67%, which might explain their increased fragmentation, reduced contig length, and inability to assign haplotigs (Table 1). The composition of TE superfamilies on primary contigs versus haplotigs was very similar (Figure S2). Both retrotransposons (Class I) and DNA transposons (Class II) cover 30% of the genome each (note that distinct TEs belonging to different categories can overlap). For Class I transposons, the long terminal repeat (LTR) order was the most prominent with ~27% coverage, and within this order elements from the Gypsy and Copia superfamilies were most prominent. The only other Class I orders with greater than 1% genome coverage were LARD and DIRS elements. Class II elements were dominated by TIR elements with a genome coverage of ~20%, with significant contributions of elements belonging to the hAT, MuDR, PIF-Harbinger, Tc1-Mariner, and CATCA superfamily. More than 6% of the genome was covered by Class II elements that could not be classified below the class level and showed no homology to previously identified TEs. This is in contrast to the minimal coverage by unclassifiable Class I elements (0.05%).

Overall, this is the highest number of identified transposable elements detected in any *Pst* genome assembly so far, as previous reports varied from 17% to 50% (4, 7, 29). Such an increased content of identified transposable elements is likely due to the increased contiguity and the absence of any unidentified bases (Ns) in our assembly (Figure 1).

Next, we reasoned that younger, less divergent TEs are mostly likely to contribute to current genome evolution. Therefore, we estimated TE age on primary contigs, which are more contiguous than haplotigs, based on their divergence from the consensus sequence of each element (Figure S3A and B, and File S1) (39). This enabled us to investigate how much of the genome is covered by relatively young TEs (< 100 Mya in our approximation) with high copy numbers (> 50 copies) (Figure S3C). The genome coverage of these younger high copy number TEs followed the overall coverage analysis closely (Figure S2B and C and S3C). Class I:LTR elements, especially Copia and Gypsy superfamily members, and Class II elements belonging to the TIR order and unclassified Class II elements are likely to contribute to current genome evolution. In future, the availability of further high quality genome assemblies for rust fungi will provide greater insight into TE evolution in *Pucciniales* and their contribution to genome evolution.

### High levels of inter-haplotype structural variation lead to variable gene content between primary contigs and haplotigs

We also annotated gene models on primary contigs and haplotigs independently using extensive sets of newly generated and publicly available RNA-seq data (40). This is in contrast to previously published *Pst* genomes that are annotated nearly exclusively using *ab initio* gene finding approaches without gene expression data (4, 5, 7, 29). The newly generated RNA-seq datasets were obtained from dormant and germinated urediniospores, wheat leaf tissue six and nine days post infection (dpi), and haustoria-enriched fractions. These datasets were complemented by publicly available RNAseq data from germinated spores, and infected wheat tissue sampled at 13 different time points x plant genotype combinations (40). We used these extensive expression data in a comprehensive genome annotation pipeline (41–45) and identified 15,928 and 14,321 gene models on primary contigs and haplotigs respectively, after filtering for genes related to TE function (Tables 1 and S2) (46, 47). The protein sequences of these genes were functionally annotated using a number of bioinformatic tools (Table S2 and File S2) (32, 48–52). We obtained very similar annotation levels for primary contigs and haplotigs with about 52% of all proteins having at least one functional annotation in the following categories; GO terms, InterPro match, Pfam domain, EggNog term, KEGG pathway annotation, MEROPs catalytic domain, or carbohydrate hydrolyzing enzymatic domains (CAZy) (32, 48–52). The level of functional annotation for *Pst* proteins identified as BUSCO orthologs was near complete with only three proteins in total (< 0.1%) lacking any functionally recognizable domain (Table S2). This pattern was reversed when characterizing candidate effectors (see identification below) as approximately 83% of all proteins lacked a conserved functional domain.

Overall, the haplotype-phased assembly did not show biased distribution of any particular gene annotation group (Table S2); this is consistent with the high level of haplotype phasing. This encouraged us to investigate the relationship between the two haplotype-phased block assemblies (primary contigs compared to haplotigs) in terms of gene content. One must keep in mind that these two assemblies do not actually represent the true haploid genomes, because of potential haplotype switching between primary contigs and haplotigs, and the inability to assign independent contigs to a specific haploid genome copy (30). However, a relational comparison between the two assemblies is still valuable in order to investigate the approximate inter-haplotype gene diversity. Therefore, to simplify the analysis we treated primary contigs and haplotigs as two representative genetic units. We used Proteinortho in synteny mode to identify allele pairs between the primary contigs and haplotigs (53). We identified a total of 10,921 potential syntenic allele pairings including 10,785 primary proteins and 10,860 haplotig proteins (Table S3, File S3 and S4 for allelic variation comparison). Of these, 9,756 were properly paired where the haplotig gene models were located on an associated haplotig that overlapped with the primary gene model when performing targeted whole-genome alignments (Figure S4A, Table S2). These correspond to ‘classic’ alleles in a diploid organism. Another 450 pairs were not directly linked as the haplotig containing the allelic ortholog did not overlap with the primary gene model although it was associated with the primary contig (Figure S4B and File S3). These may be simple rearrangements linked to inversions or repeat duplications. A further 715 pairs were completely unlinked as the allele-containing haplotig was not associated with the respective primary contig in our assembly (Figure S4C and File S4). We randomly selected 176 of these loci and investigated them manually by whole genome alignment of haplotigs to primary contigs, followed by micro-synteny analysis of the identified gene loci (35, 54, 55). An example of this analysis is illustrated in Figure 3. In this case, a ~40 kb region present both in primary contig 014 and haplotig 027_006 showed micro-synteny for three genes each, namely *Pst104E_05635-05637* and *Pst_104E_24450-24452*, respectively (Figure 3D), while the overall macro-synteny was not conserved (Figure 3A-C). This may have been caused by genetic transposition of the identified region from the chromosomal region corresponding to a haplotig that fully aligned with primary contig 014 into the sequence of the chromosomal region corresponding to haplotig 027_006. We found support for such allele transposition, either via cut and paste or copy and paste mechanisms, in 71/176 cases. The remaining cases could not be categorized confidently and may represent complex genomic regions, genetically linked contigs that were broken up during the assembly process, gene duplication events, or miss-assemblies. Based on this manual inspection we estimate that approximately 280 loci (71/176*715 total pairs) contain alleles that might be rearranged in one of the two haploid genomes. We identified a further 912 loci that clustered at the protein level yet their genomic location was not syntenic between the two haplotype-phased block assemblies (File S5). We refer to these genes as inter-haplotype paralogs. In summary, this suggests that over 3% (~1,192/30,249) of all genes are closely related at the protein level but do not reside in regions displaying macro-synteny.

**Figure 3:**
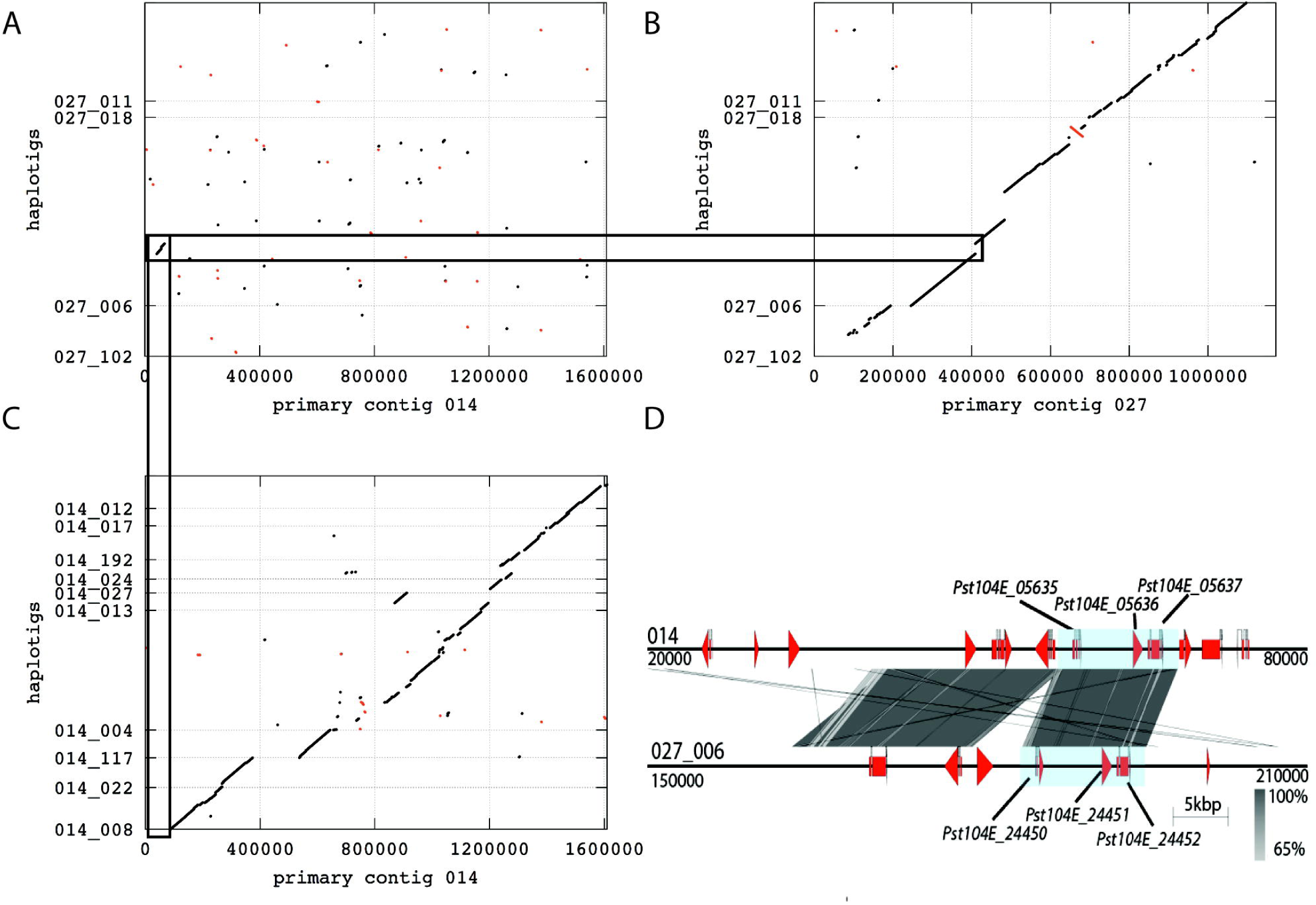
Allele transposition in the Pst-104E genome. A-C) Dot plots of whole genome alignments generated using the mummer toolset where the x-axis represents primary contig and the y-axis the haplotig sequence. A) shows the whole genome alignments of haplotigs_027_xxx to primary contig 014. B) shows the whole genome alignment of haplotigs_027_xxx to primary contig 027. C) shows the whole genome alignment of haplotigs_014_xxx to primary contig 014. Black lines indicate alignments in the forward direction and red lines in the reverse direction in the haplotig sequence. The black rectangles highlight a ~40 kb region in haplotig_027_006 that does not align to primary contig 027 yet aligns to a region in primary contig 014, which is not covered by an associated haplotig of 014. In D) we show micro-synteny analysis of this extended region with primary contig 014 on top and haplotig_027_006 on the bottom. Gene models identified as alleles are labeled with their locus tag and shaded by a light blue background. Vertical grey shading illustrates the blastn identity between sequences on both contigs according to the scale shown in the right bottom corner next to the sequence scale bar. Start and stop positions for each contig sequence are given at the start and the end of each contig.

We identified 4,761 primary and 2,931 haplotig genes that did not cluster at the protein level using Proteinortho and hence may represent singletons, with singletons defined as genes of a diploid/dihaploid organism that lack alleles or inter-haplotype paralogs (Table S3). Of the 4,761 primary genes, 663 were located in regions where the assembly was not haplotype-phased based on coverage analysis using Illumina short-read data (File S6). Hence we identified 7,029 true singletons (File S7) when comparing both haplotype-phase block assemblies, and 1,506 of these singletons are referred to as single haplotype genes (File S8) because they lacked any BLAST hit (blastn, e-value < 0.01) when using the gene sequence as a query against the alternate haplotype-phase block sequence. These single haplotype genes are often linked in clusters, because for 1,164 single haplotype genes at least one of their nearest neighbors is also a haplotype-specific gene, compared to 212 of an equally sized random subsample of all genes (Fisher’s exact test, p-value ~ 2.3*10^−109^). Similarly, 1,492 haplotype-specific genes are located in regions where primary contigs and associated haplotigs do not align, indicating haplotype specific regions. Single haplotype genes are highly enriched in these regions as only 251 of an equally sized random subsample of all genes displayed a similar location (Fisher’s exact test, p-value ~ 4.5*10^−265^). Taken together, these findings suggest that there are numerous large presence-absence structural polymorphisms between the two haploid genomes that can span multiple adjacent genes, and therefore contain many of the haplotype-specific genes. To study the overall conservation of these single haplotype genes we queried them against the EnsemblFungi cDNA and NCBI nr databases (blastn, e-value < 0.01) (56, 57). Out of 1506 genes, 1424 had at least one significant hit in either database, with the top hits in all cases being fungal sequences. The remaining 82 genes lacked any sequence homology to known fungal genes. These genes were significantly shorter compared to all genes (mean length 538 bases versus 1538, two-sided Student’s t-test, p-value ~ 2.38e^−07^). We identified expression evidence for 27/82 of these genes including 7 of 10 predicted candidate effectors. This is consistent with observations in other fungi for which isolate-specific genes tend to be shorter and are lower expressed than genes that are conserved between isolates (58). Overall, the high levels of non-allelic genes (~25%) and single haplotype genes (~5%) illustrates that the large inter-haplotype polymorphism at the nucleotide and structural levels (Figure 2 and 3B, C) results in significant differences in gene content.

### Candidate effector gene prediction using machine learning and *in planta* expression data

The diversity of plant pathogen effectors makes them impossible to identify based on protein sequences alone (59). Only a small number of effectors have thus far been confirmed in rust fungi, namely AvrP123, AvrP4, AvrL567, AvrM, RTP1, PGTAUSPE-10-1 (60), AvrL2 and AvrM14 (61), PstSCR1 (62) and PEC6 (63). At the sequence level, effectors do not share common domains or motifs, apart from the presence of a signal peptide. To predict candidate effectors in *Pst*-104E, we utilized a combination of gene expression analysis and machine learning methods. First, we predicted fungal rust secretomes based on a protocol optimized for recovering fungal candidate effectors (64). We observed large differences in secretome sizes across rust proteomes, e.g. the stripe rust isolate *Pst*-887 had a small secretome compared to Pst-104E (Table S4). Overall the number of secreted proteins appeared to correlate with completeness of *Pst* genome assemblies based on BUSCO analysis (Figure 1B and Table S4). This implies that it is difficult to perform comprehensive orthology analyses between current *Pst* assemblies given that many appear to be incomplete in terms of BUSCOs and therefore are likely incomplete for other gene families also, including secreted proteins.

To predict candidate effectors, we used the machine learning approach EffectorP on all secreted proteins without predicted transmembrane domains (64). Overall we identified 1,069 and 969 candidate effectors from primary contigs and haplotigs respectively (File S9). We complemented this *in silico* approach with a detailed expression analysis of Pst-104E genes that encode secreted proteins. We used gene expression data and *k*-means clustering to predict clusters in the secretome that are differentially expressed during infection and exhibit similar expression profiles (Figure 4, File S10). For the primary contigs of *Pst*-104E, this resulted in eight predicted clusters. The expression profiles of three clusters (clusters 2, 3 and 8) resembled the expected expression patterns of haustorially-delivered cytoplasmic rust effectors, namely a high expression in haustorial tissue and at the infection time points (6 and 9 dpi) as well as low expression in spores (Figure 4A). In total there are 809 genes in clusters 2, 3, and 8, of which 306 (~38%) were also identified by EffectorP as candidate effectors (Table S5). Upon closer inspection of primary contig expression patterns, cluster 8 in particular exhibits the highest overall haustorial expression and overall lowest expression in spores, indicating it is likely to contain cytoplasmic effectors. Interestingly, whilst cluster 8 shows the lowest percentage of EffectorP predicted candidate effectors (26%), it has the highest percentage of proteins with a predicted nuclear localization signal (NLS) (Table S5) (65). We also observed that proteins in cluster 8 are mostly larger (average length of 410 aa) than other known rust effectors (the largest is AvrM, at 314 aa), which might indicate that *Pst* utilizes a class of larger effector proteins that target host nuclei. Similarly, oomycete pathogens secrete a class of cytoplasmic effectors called Crinklers that carry NLSs (66, 67) but these are not predicted as candidate effectors by EffectorP, possibly due to their larger size. Therefore, we included both *in planta* up-regulated secreted proteins as well as EffectorP predicted proteins as candidate effectors. In total, we identified 1,572 candidate effectors on primary contigs when combining predictions based on *in planta* expression analysis and EffectorP. We identified similar expression patterns for secreted proteins on haplotigs. Clusters 11, 13, 14 and 15 shared a similar expression profile to clusters 2, 3, and 8 and contained 673 genes (Table S6 and S7). Of these, 234 (~37%) were also identified by EffectorP amounting to a total of 1,388 candidate effectors on haplotigs. Overall, we identified a set of 1,725 non-redundant candidate effectors, identified by machine learning and expression analysis approaches, when combining all candidate effectors on primary contigs and haplotigs (File S11).

**Figure 4:**
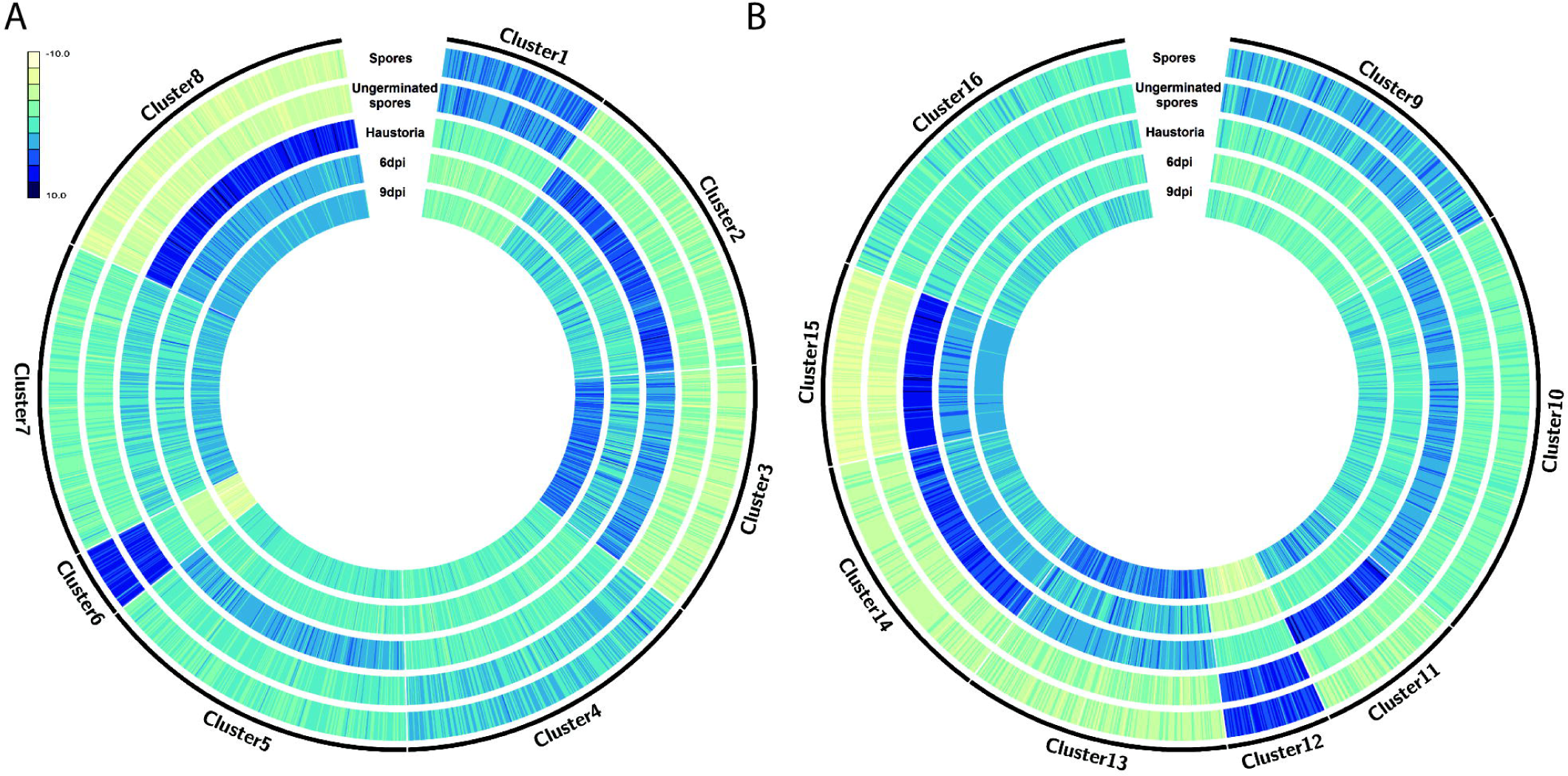
Identification of candidate effectors based on detailed expression analysis of secreted proteins of both Pst-104E assemblies. A) Clustering of Pst-104E secretome expression profiles for genes located on primary contigs. Blue color intensity indicates the relative expression level using rlog transformed read counts in spores, germinated spores, haustoria, and in wheat tissue at 6 and 9 days post infection. For example, cluster 8 shows the lowest relative expression in spores and the highest in haustoria compared to the other clusters. B) Clustering of Pst-104E secretome expression profiles for genes located on haplotigs.

### Candidate effector genes are spatially associated with conserved genes and with each other

For many filamentous plant pathogens a ‘two speed’ genome has been suggested to contribute to rapid evolution in terms of candidate effector variability (68). For example, in fungal plant pathogens such as *Fusarium oxysporum* spp. and *Verticillium dahliae*, lineage specific genomic regions and/or dispensable chromosomes are enriched for TEs and candidate effector genes. In several *Phytophthora* spp., candidate effectors have been reported to localize in gene sparse, TE-rich regions, which show signs of accelerated evolution (68, 69, 74). It is not known if rust genomes have a comparable genome architecture that facilitates rapid evolution of candidate effector genes. Hence we investigated the genomic location of candidate effectors in relation to several genomic features including TEs, neighboring genes, BUSCOs, other candidate effectors, and AT content (Figures 5, 6, S5, and S6). We focused mostly on candidate effectors on primary contigs, because the primary assembly is far more contiguous compared to its haplotigs thereby facilitating our analysis (Figure 1). In addition, we made use of our haplotype-phased assembly and investigated if allelic candidate effector variants show distinct features when compared to haplotype singletons. In all cases we used a random subset of genes and BUSCO gene sets as control groups. We envisioned BUSCO genes as a particularly well-suited control group as these are conserved within the phylum of basidiomycetes (75) and can therefore be considered as part of the *Pst* core genome. On the contrary, candidate effector genes are reported to be more specific to the class, species, or isolate level (6, 76). This observation also holds true for *Pst*-104E, because we only observed 40 BLAST hits outside the class of *Pucciniomycetes* for 1,725 non-redundant candidate effectors using EnsemblFungi cDNA as reference (blastn, e-value 1*e^−5^).

**Figure 5:**
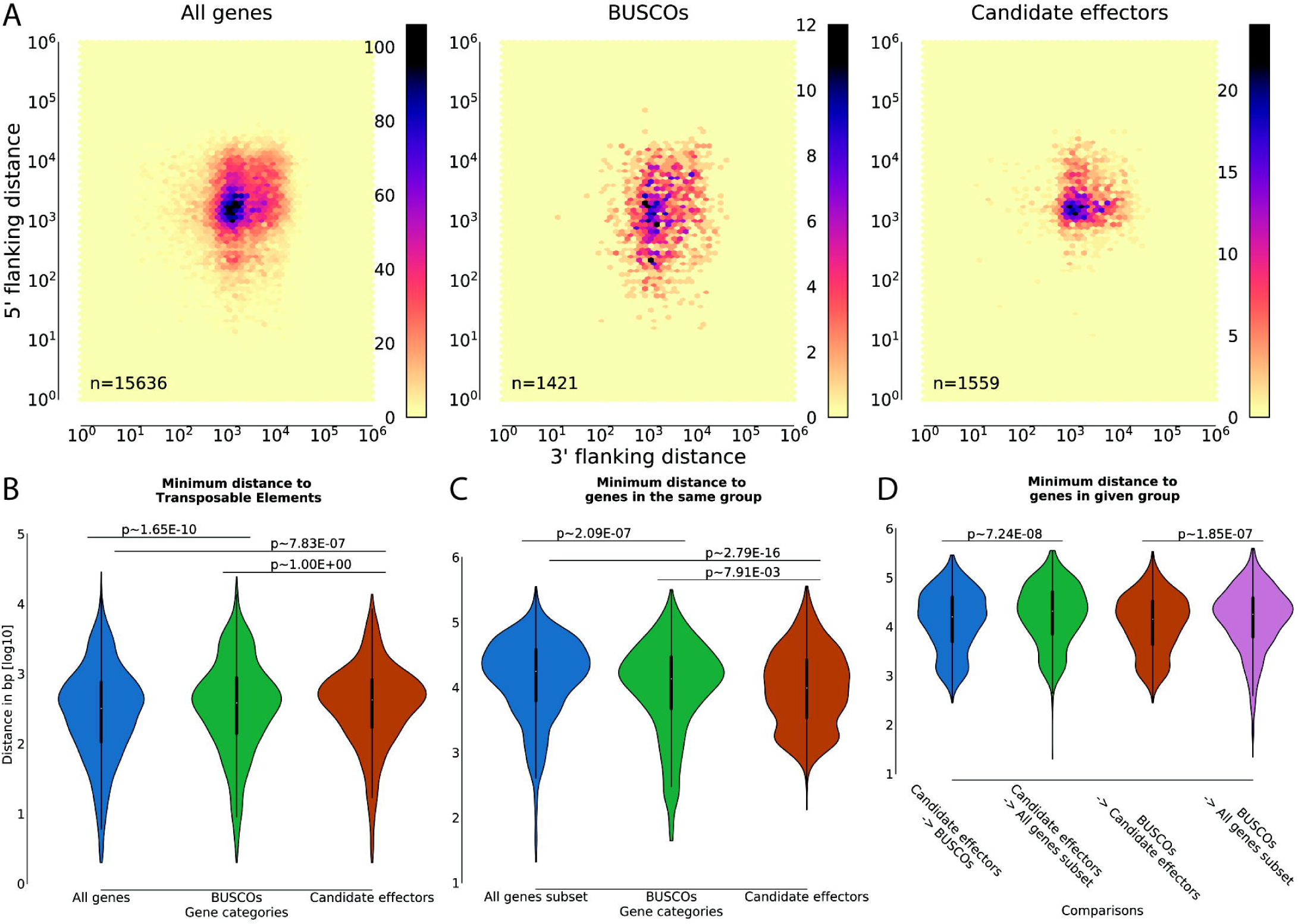
Candidate effector genes are spatially associated with conserved genes and with each other. A) Nearest neighbor gene distance density hexplots for three gene categories, including all genes, BUSCOs and candidate effectors. Each subplot represents a distance density hexplot with the log10 3’-flanking and 5’-flanking distance to the nearest neighboring gene plotted along the x-axis and y-axis, respectively. B) Violin plots for the log10 distance to the most proximal transposable element for genes in each category without allowing for overlap. C) Violin plots for the log10 distance to the most proximal gene in the same category for subsamples of each category equivalent to the smallest category size (n=1444). D) Violin plots for the minimum distance [log10] of candidate effectors and BUSCOs to each other or a random subset of genes (n = 1444). The p-values in B, C, and D are calculated using the Wilcoxon rank-sum test corrected for multiple testing (Bonferroni, alpha=0.05) on the linear distance in bases.

**Figure 6:**
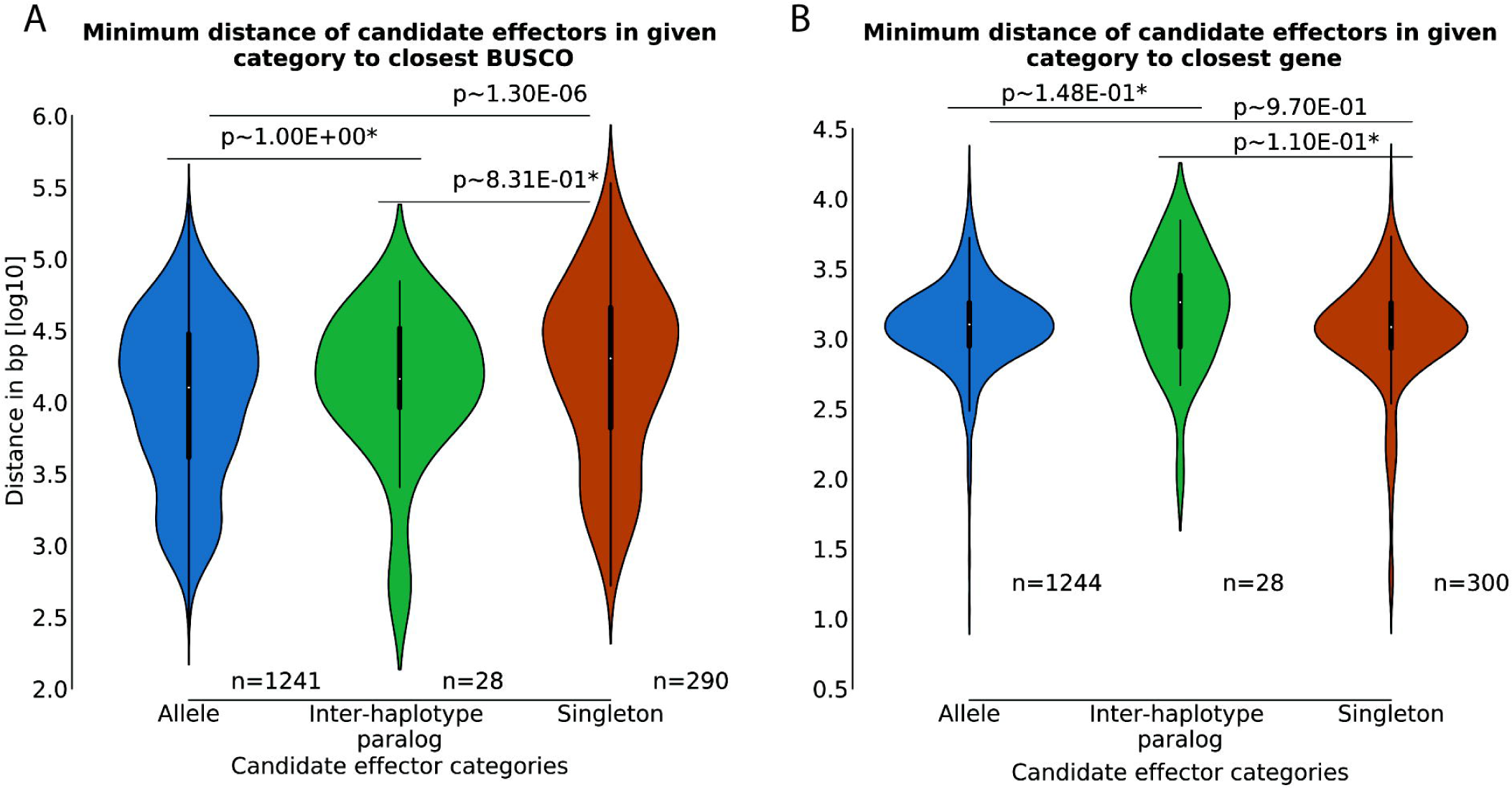
The candidate effector allele status influences association with conserved genes. A) Violin plots for the log10 distance to the most proximal BUSCO for candidate effectors in each category. The Kruskal–Wallis one-way analysis of variance of all three categories shows a significant difference between the three samples (p ~ 2.36e^−06^). B) Violin plots for the log10 distance to the most proximal gene for candidate effectors in each category. The Kruskal–Wallis one-way analysis of variance of all three categories shows no significant difference between the three samples (p ~ 0.08). The p-values in A and B are calculated using the Wilcoxon rank-sum test corrected for multiple testing (Bonferroni, alpha=0.05) on the linear distance in bases. *Wilcoxon rank-sum test comparisons with inter-haploid genome paralogs lack statistical power due to the small samples size of n=28.

We first tested if candidate effectors are located in gene sparse regions when compared to all genes or BUSCOs. For this analysis, we generated density plots using the distances from the 5’ and 3’ ends of each gene to its closest neighbor in either direction (68). When comparing gene distance density hexplots we observed very similar distributions between candidate effectors and all genes. Candidate effectors in general did not appear to be located in gene sparse regions and neither did BUSCOs (Figure 5A). Similar effects have been reported for other rust species, such as the oat crown rust pathogen *Puccinia coronata* f. sp. *avenae* (77). Next, we tested if candidate effectors are linked to TEs as observed for other plant pathogenic fungi (74). We compared the minimum distance of all genes, BUSCOs and candidate effectors to TEs. Candidate effectors globally did not display a preferential association with TEs when compared with genes in general (Figure 5B). However, on close examination of the relative spatial distribution of TEs, candidate effectors and BUSCOs on the 30 largest contigs we could identify some regions where candidate effectors are closely associated with TEs (Figure S5). The observation that candidate effectors are not associated globally with TEs is consistent with reports of other rust fungi including *P. coronata* f. sp. *avenae, P. graminis tritici*, and *Melampsora larici-populina* (6, 77). In the case of *Pst*, we aim to address the question of the involvement of TEs in the evolution of novel virulences by re-sequencing Pst-104E mutant progeny with distinct virulence profiles collected in Australia between 1980 and 2003 (27).

The observation that candidate effectors and BUSCOs show similar localization patterns relative to all genes and TEs led us to investigate if these two gene groups are spatially associated, and if each group clusters with itself. We first compared the minimum distance between genes of the same group when subsampling to an equal number of genes in each group. Indeed, when comparing the minimum distances between candidate effectors we found that these were less than the minimum distances between a random subset of genes (Figure 5C). BUSCOs were also more closely associated with each other than a random subset of genes. Consistently, when we investigated the number of candidate effectors that clustered within a minimum given distance we found that they are more clustered than BUSCOs or an equal sized random subset of all genes (Figure S6). A similar trend yet to a lesser degree was observed for BUSCOs. Clustering of candidate effectors was also identified as a feature of several smut fungi including *Ustilago maydis* and *Sporisorium scitamineum* (3, 78). In these related basidomycete plant pathogens, candidate effector gene clusters are born via tandem duplication and linked TEs are hypothesized to contribute to the rapid evolution of these genes.

The observed spatial association of both BUSCOs and candidate effectors with themselves led us to investigate if these two gene groups are spatially associated with each other. Indeed, candidate effectors were located more closely with BUSCOs and vice versa when compared to a random subsample of all genes (Figure 5D). This is a surprising observation because BUSCOs are defined by their overall conservation while candidate effectors are far less conserved. In obligate biotrophic fungi, a subset of effectors may be essential, because host colonization is an absolute requirement for survival. Therefore, there may be selection pressure on obligate biotrophs to favor recombination events that link some essential effectors to other essential genes (e.g. BUSCOs) to ensure their inheritance and conservation within the species complex. This is in contrast to plant pathogens that are also able to grow saprophytically such as *Z. tritici, V. dahliae, U. maydis* and *P. infestans* (3, 72, 73, 79). In addition, the genetic variation within *Pst* isolates in its center of genetic diversity is high, and sexual recombination may generate diverse effector complements that allow colonization of taxonomically distinct hosts including barberry and grasses. In these natural environments, the composition of effector complements may be selectively neutral and these processes may not facilitate effector gene compartmentalization. Once *Pst* leaves the Himalayan region and invades large wheat growing areas, sexual recombination is absent and hence effector gene compartmentalization is not possible.

### The candidate effector allele status influences association with conserved genes and evolutionary conservation

We next investigated if the distance between candidate effectors and BUSCOs is correlated with their allelic variation. We calculated the normalized Levenshtein distance of cDNA and amino acid alignments for all allele pairs. The normalized Levenshtein distance measures the required single-character edits (insertions, deletions or substitutions) to convert two strings into each other, e.g. an alignment of two allele sequences, while accounting for differences in sequence length. It can therefore be used as a proxy for sequence variation between two alleles (80). We did not observe any significant difference between the Levenshtein distance at the cDNA level when comparing BUSCOs and candidate effectors, whereas alleles of all other genes were more variable than candidate effectors (Table 2). This was in contrast to the variation seen at the protein level, where candidate effectors were more variable than BUSCOs (Table 2). This suggests that for candidate effectors, changes at the DNA level are more likely to result in changes to the protein sequence. We therefore also calculated the ratio of nonsynonymous to synonymous mutations for all alleles (dN/dS ratio) wherever possible (81). Indeed, analysis of the dN/dS ratios supported our previous observation that for candidate effectors, changes in the DNA sequence were more likely to alter the protein sequence (Table 2). This suggests that candidate effectors evolve faster than BUSCOs and most other allele pairs even though they are spatially associated with BUSCOs. The sequence variation in candidate effector allele pairs was not correlated with distance to the closest BUSCO, using either Levenshtein distances on the protein level or dN/dS as a proxy (Spearman, correlation < |0.06|, p > 0.15). Subsequently, we investigated if candidate effector singletons were more distant from BUSCOs than their paired-allele counterparts. These singletons have either diverged dramatically from their ancestral allele counterparts, were lost due to structural rearrangements and mutations, or encode *de novo* evolved candidate effectors. The candidate effector singletons were found to be located more distantly from BUSCOs than paired-allele candidate effectors (Figure 6A), but were not more distant from other genes in general (Figure 6B). Nonetheless, we reasoned that these candidate effector singletons might be more likely to be isolate- or species-specific given their distinct genomic locations compared to paired-allele candidate effectors. We tested if candidate effector singletons are more likely to lack orthologs in publicly available *Pst* genomes or other genomes of *Pucciniales* species (82). Out of a total of 453 candidate effector singletons, 116 lacked an ortholog in five other *Pst* genomes, compared to 118 out of 1,272 allelic candidate effectors. Singletons are therefore more likely to be isolate-specific than are paired-allele candidate effectors (Fischer’s exact test, p ~ 1.36e^−16^). We observed a similar trend when comparing Pst-104E with the six publically available *Pucciniales* genomes. Of 985 candidate effectors lacking orthologs in other rust fungi, 313 were singletons and 672 allelic, also showing a enrichment for candidate effector singletons (Fischer’s exact test, p ~ 4.45e^−26^).

**Table 2:** Candidate effector alleles are more variable than BUSCOs alleles on the protein level Summary of normalized Levenshtein distances and dN/dS ratios calculated for CDS alignments and codon based amino acid sequence alignments. A Percentage of genes or proteins for which the normalized Levenshtein distance is greater 0. B are calculated using the Wilcoxon rank-sum test corrected for multiple testing (Bonferroni, alpha=0.05). C Number of loci for which dN/dS ratios could be calculated using yn (ref.). D Percentage of loci for which dN/dS was not 0.

|  | BUSCOs | Candidate<br>effectors | Other Genes |
| --- | --- | --- | --- |
| #of loci with Levenshtein distance<br>CDS | 1198 | 1214 | 8509 |
| % of genes showing variation<sup>A</sup> | 81 | 66 | 79 |
| Median | 0.0069 | 0.0044 | 0.0074 |
| Mean | 0.0288 | 0.0409 | 0.0579 |
| Wilcoxon-rank Sum test vs.<br>candidate effectors<sup>B</sup> | ~9.47*e-02 | N.A. | ~1.21*e-10 |
| #of loci with Levenshtein distance<br>protein | 1198 | 1214 | 8509 |
| % of proteins showing variation<sup>A</sup> | 65 | 60 | 70 |
| Median | 0.0028 | 0.0060 | 0.0075 |
| Mean | 0.0264 | 0.0474 | 0.0637 |
| Wilcoxon-rank Sum test vs.<br>candidate effectors<sup>B</sup> | ~9.46*e-05 | N.A. | ~1.86*e-10 |
| #of loci with dN/dS ratio<sup>C</sup> | 859 | 619 | 5403 |
| %of loci showing variation<sup>D</sup> | 75 | 87 | 85 |
| Median | 0.0802 | 0.3972 | 0.2840 |
| Mean | 0.2012 | 0.4797 | 0.3432 |
| Wilcoxon-rank Sum test vs.<br>candidate effectors<sup>B</sup> | ~2.42*e-54 | N.A. | ~1.91*e-06 |

### Conclusions

Using long-read sequencing technology we are now starting to uncover the genomic diversity of dikaryotic fungi that was previously hidden by a reliance on short-read sequence assemblies. We used this approach to generate a highly contiguous haplotype-phased assembly of the Australian founder *Pst* pathotype. We are now able to describe the levels of inter-haplotype diversity both on the structural and gene levels. It is difficult to fully evaluate the significance of observed levels of variations without additional experiments and in the absence of similar studies. With over 6% variation, the inter-haplotype diversity of Pst-104E is higher than that reported for *P. coronata* f. sp. *avenae* which ranges between 2.1 and 2.7% (77). It is also higher than the variation observed between two isolates of *Z. tritici* (3D7 vs. MG2, 4.9%), an ascomycete pathogen of wheat that undergoes frequent sexual cycles (58, 72), and two isolates of *V. dahliea* (JR2 vs. VdLs17, 1.7%), an ascomycete pathogen of tomato that propagates almost exclusively asexually (73). These comparisons suggest that the observed inter-haplotype diversity of *Pst* is high. Pst-104E belongs to the ‘North Western European’ (NW European) lineage of *Pst,* which has undergone long-term asexual reproduction. The NW European *Pst* lineage can be traced back to its first sampling in mid-1950 in the Netherlands, and has not shown any signs of sexual recombination since (21, 83, 84). Consistent with this, two *Pca* isolates that show much less inter-haplotype variation than Pst-104E are from populations that reproduce both sexually and asexually on common buckthorn and oat, respectively (77). Frequent sexual recombination is likely to reduce inter-haplotype diversity and to purge mutations that are deleterious in the monokaryon stage (85). On the other hand, long-term clonal lineages might accumulate polymorphisms that clear unwanted *Avr* genes but also contribute to genomic decay. It has long been hypothesized that prolonged clonal reproduction in the absence of sexual recombination and chromosomal re-assortment will lead to high levels of heterozygosity between chromosomes that were initially homologous, a phenomenon known as the Meselson effect (86). This also suggests that *Pst* isolates from the center of genetic diversity may display less inter-haplotype diversity and a reduced allelic variation due to sexual recombination. This is an aspect of *Pst* biology that we are aiming to test in future studies. With respect to this, it is an interesting point if *Pst*-104E is still viable as a monokaryon in the absence of selection to retain gene function related to infection of barberry. The accumulation of large scale polymorphisms and potentially deleterious mutations in each haploid genome of *Pst*-104E might have been buffered in the dikaryon stage, but it is likely that it represents a terminal lineage of *Pst* in agreement with Muller’s rachet hypothesis (85). Isolates from the NW European lineage show a reduction in teliospore production on wheat, the entry point into the *Pst* sexual cycle, when compared to isolates from the Himalayan region where sexual reproduction is common (87). Also, successful sexual reproduction under laboratory conditions has been reported only for *Pst* isolates that emerged recently from the center of diversity in the Himalayan region (88), but not for isolates that have undergone long term clonal reproduction such as the NW European lineage (personal communication J. Rodriguez-Algaba). Lastly, *Pst* populations of the NW European lineage have been completely replaced by more recent *Pst* incursions in Europe and Australia (15, 25).

In future it will be important to generate high quality genomes for more *Pst* isolates, including from sexual populations in the Himalayan regions (89). This will enable us understand the role of sexual and asexual reproduction in the genome evolution of a dikaryon in the wild versus agricultural settings. For now, the near-complete haplotype-phased genome of *Pst*-104E provides a first haplotype-aware insight into the genetic architecture of a dikaryotic rust fungus pathogenic on wheat. In itself it is a high quality reference genome enabling investigation of the rapid and devastating evolution of the fungus to virulence during its asexual reproduction cycle in all wheat growing areas today.

## Material and Methods

### Data availability

The following are the NCBI accession numbers for data generated in course of this manuscript which is registered as BioProject PRJNA396589.

Short read archive accession numbers are as follows:

SRX311905-14 and SRX311918-20: PacBio 10-20kb BluePippin kit, RSII, 13 SMRT cells.

SRX311916 and 17: genomic DNA TruSeq library, HiSeq 2000, 100 bases paired end library.

SRX311915: genomic DNA TruSeq PCR free, MiSeq, 250 bases paired end library.

SRX3191029-43: TruSeq v2 RNAseq samples, HiSeq 2000 100 bases paired end library.

Bioinformatic scripts, supplemental files and genome annotation can be found on this manuscripts github page https://github.com/BenjaminSchwessinger/Pst104E137A-genome.

The genome is also available at MycoCosm (https://genome.jgi.doe.gov/Pucstr1/Pucstr1.home.html).

### *Puccinia striiformis* f. sp. *tritici* pathotype, growth conditions and spore amplification

The isolate of pathotype 104E137A- was collected from the field in 1982 (Plant Breeding Institute accession 821559=415), tested and propagated as described previously (27). This pathotype is virulent on Heines VII (*Yr2*, Yr25), Vilmorin 23 (*Yr3*), Hybrid 46 (*Yr4*), Stubes Dickkopf, Nord Deprez, Suwon92/Omar and Avocet S (27). The rust propagated for PacBio sequencing was produced by selecting a single pustule of the original isolate (increase 0415Ga) on wheat plants of the susceptible variety ‘Morocco’. The initial inoculation involved rubbing leaves of the susceptible host with a spores from a sterile cotton tip. Plants were incubated under plastic in the dark at 9.5°C for 18 h before being transferred to a greenhouse microclimate set at 22°C ±2°C. After 6 d, plants were observed and all leaves were removed except for one leaf which showed signs of infection by a single fleck indicating a rust pustule was soon to erupt from the location. After pustule eruption, the single pustule selection was repeated to ensure that the starter material for propagation was a single genotype. Multiplication of rust was done on *Triticum aestivum* cv. ‘Morocco’. For multiplication, 20 seeds of ‘Morocco’ were placed as a single layer into four inch pots filled with pasteurised soil and watered with a half-strength solution of liquid fertiliser (Aquasol, Yates). At full coleoptile emergence, each pot was treated with 50 mL maleic hydrazide solution (2 mL L^−1^ Slow Grow 270, Kendron). At full leaf emergence plants were inoculated by rubbing with the pustules formed in the previous step and incubated as described previously. Once four pots of ‘Morocco’ were heavily infected, spores were collected and inoculated onto 64 four inch pots and a differential set to check pathotype identity and purity. Rust spores were collected from the 64 pots using a GRA-101 large spore cyclone (Tallgrass Solutions) attached to a domestic vacuum cleaner. Spores were dried over silica gel for 7 days before being sieved through a 50 μm sieve and being stored at -80 °C until DNA extraction.

### DNA extraction and genome sequencing

DNA was extracted from dried dormant *Pst* urediniospores as described it in detail elsewhere (90, 91). PacBio sequencing was performed at the Ramaciotti Centre (Sydney, Australia). For library preparation the 20 kb BluePippin kit (PacBio) was used. DNA libraries were sequenced on a PacBio RSII instrument using P6-C4 chemistry. In total we sequenced 13 SMRT cells (Table S1). DNA samples from the same *Pst* pathotype were also sequenced with Illumina short read technology. We sequenced one TruSeq library on a HiSeq 2000 instrument as a 100 bases paired end library at the University of Western Sydney (Sydney, Australia). We sequenced one TruSeq PCR free 250 bases paired end library on an Illumina MiSeq instrument at the Ramaciotti Center (Sydney, Australia).

### Genome assembly and manual curation

For genome assembly we used FALCON-Unzip github tag 1.7.4 with the parameters described in File S12 and S13 (30). We checked the resulting contigs for eukaryotic contamination by blastn searches against the NCBI nucleotide reference database (downloaded 04052016) (38). None of the contigs had predominant non-eukaryotic sequences as best BLAST hits at any given position. We performed two manual curation steps. In the first step we reasoned that some of the primary contigs without haplotigs may actually represent haplotigs that could not be connected to their respective primary contigs in the assembly graph because of too large a difference between the two haplotypes. We aligned all primary contigs without haplotigs to primary contigs with haplotigs using mummer version 3 (35). We screened the best alignments of each primary contig without haplotig for percentage alignment, length of alignments, and if they align to regions in the primary contigs that previously had not been covered by a haplotig alignment. Using this approach we re-assigned 55 primary contigs without haplotigs (~6 Mb) to haplotigs (Table S8). In the second step of manual curation we removed all contigs with an mean coverage of greater 2000x when using Illumina short read data. In total we removed 18 primary contigs (~0.6Mb) and 7 haplotigs (~0.2Mb) of which most were mitochondrial contigs based on blastn analysis. The final assembly contains 156 primary contigs (~83Mb) and 475 haplotigs (~73Mb) (Table S8).

### Coverage analysis and identification of unphased regions in primary contigs

We aimed to assess the coverage within contigs and between contigs when mapping Illumina short read data on primary contigs (p) and primary contigs and haplotigs (ph) at the same time. We reasoned that unphased region of primary contigs should have about twice the coverage of phased regions when mapping against ph and similar coverage comparing mapping against p vs. ph. We trimmed Illumina short reads using Trimmomatic v0.35 (92) (ILLUMINACLIP:adapter.fa:2:30:10 LEADING:3 TRAILING:3 SLIDINGWINDOW:4:25 MINLEN:35) and assessed read quality with FastQC v0.11.4 (93). Reads were mapped against primary contigs only or primary contigs and haplotigs using BWA-MEM v0.7.15-r1142-dirty using the standard parameters (94). The coverage for each position was calculated with samtools v1.3.1 using depth with the ‘-aa’ flag (95). Unphased regions on primary contigs were defined as outlined above and converted to bed format. See jupyter notebook Pst_104E_v12_coverage_analysis_submission_21092017 in the github repo.

We also performed a detailed coverage sequence depth analysis on 1 kb sliding windows using 200 base intervals. We generated corresponding bed files with the window function in pybedtools for primary contigs and haplotigs. In addition, we generated corresponding sliding window bed files for primary contig regions that aligned with haplotig regions and for regions that lacked an associated haplotig. For this purpose, we combined initial sliding window bed files (see above) with gff files illustrating primary contig region that aligned with haplotigs (96, 97). The later gff files are based on Assemblytics alignments of haplotigs to their respective primary contigs using nucmer (34). These bed files where used to calculate the mean base sequence depth based on the samtools function bedcov (95). For details on how we generated the Assemblytics based gff file see

Pst_104E_v12_defining_alleles_submission_21092017.ipynb. For details on this part of the coverage analysis see Revision_coverage_analysis.ipynb.

### Repeat annotation

Repeat regions of the primary contigs and haplotigs were predicted independently. We used the REPET pipeline v2.5 (36, 98) for repeat annotation in combination with Repbase v21.05 (46). First, we used TEdenovo to predicted novel repetitive elements following the instructions (99) using the parameters given in File S14. The set of TEs provided by Tedenovo were used to annotate all repetitive elements using Teanno following the instructions including the methodological advice (100) using the parameters given in File S15. Annotation was performed on genome version 0.4 and subsequently filtered for version 1.0 (Table S7). We transferred the superfamily annotation according to Wicker (37) for all elements from the underlying database hits if these agreed with each other and the REPET annotation. See jupyter notebooks Pst_104E_v12_TE_filtering_and_summary_p_contigs_ submission_21092017 and Pst_104E_v12_TE_filtering_and_summary_h_contigs_ submission_21092017 in the github repo for full analysis details.

### Estimation of TE age

We estimated TE age based on the divergence of each sequence identity from the consensus sequence (39). We calculated mean percentage identity for all identified TEs (repbase2005_aaSeq, repbase2005_ntSeq, and de novo identified repeats using TEdenovo) using the REPET pipeline function PostAnalyzeTELib.py -a 3 (File S1). We used the function T=D/t to roughly approximate TE age. T is the elapsed time since the ancestral sequence, D is the estimated divergence based on percentage identity calculated by the REPET pipeline (D=1-meanPctID/100), and t is the substitution rate per site per year. We estimated t~2*10̂-9 based on previous publications (101, 102). For details see notebook Revision_TE_filtering_and_summary_p_contigs.ipynb.

### Gene model annotation

We annotated genes on primary contigs and haplotigs independently. We combined RNAseq-guided *ab initio* prediction using CodingQuarry v2.0 (42) and BRAKER v1.9 (43) with *de novo* transcriptome assembly approaches using Trinity v2.2.0 (103) and PASA v2.0.1 (41). Gene models were unified using EvidenceModeler v1.1.1 (41) using the weights given in File S16.

We mapped the trimmed RNAseq reads described in this study (see below) and previously (40) against primary contigs and haplotigs using hisat2 v2.1.0 (--max-intronlen 10000 --min-intronlen 20 --dta-cufflinks) (45). For *ab initio* predictions we reconstructed transcripts using stringtie v1.2.3 (-f 0.2) (104). We ran CodingQuarry (-d) in the pathogen mode using SignalP4 (105) for secretome prediction on the soft-masked genome using RepeatMasker v4.0.5 (-xsmall -s -GC 43). Similarly, we used the stringtie reconstructed transcripts as training set for the *ab initio* prediction pipeline BRAKER 1 v1.9 (43) and used the non-repeat masked genome as reference.

We used Trinity v2.2.0 to obtain *Pst* transcripts both in the *de novo* mode and the genome-guided mode (103). Several RNAseq samples contained host and pathogen RNA as they were prepared from infected wheat tissue. We first mapped all reads to primary contigs and haplotigs using hisat2 (see above). We extracted mapped RNAseq reads using piccard tools SamToFastq (106). Only these reads mapping against *Pst* contigs were used in the *de novo* pipeline of Trinity (--seqType fq). For genome guided assembly we used bam files generated with hisat2 as starting point for Trinity (--jacard_clip, --genome_gudied_max_intron 10000). We used the PASA pipeline v2.0.2 to align both sets of Trinity transcripts against *Pst* contigs with BLAT and GMAP using the parameters given in File S17 (41).

The different gene models were combined using EvidenceModeler v.1.1.1 to get the initial gene sets for primary contigs and haplotigs (41). These were filtered for homology with proteins encoded in transposable elements. We used blastp to search for homology in the Repbase v21.07 peptides database with an e-value cut-off of 1*e^−10^. In addition, we used transposonPSI to filter out genes related to TE translocation (47, 107). We used the outer union of both approaches to remove genes coding for proteins associated with transposable elements from our list of gene models.

### Protein annotation

For initial protein annotation we used the fungal centric annotation pipeline funannotate v0.3.10 (108). This included annotation for proteins with homology to swissprot (uniref90, downloaded 22/09/2016) (50), to carbohydrate-active enzyme (dbCAN, downloaded 22/09/2016) (49), to peptidases (MEROPS v10.0) (52, 109), for proteins with eggnog terms (eggnog v4.5) (110) and SignalP4 (105). This annotation was complemented by interproscan v5.21-60 (-iprlookup -goterms -pa) (48), eggnog-mapper v0.99.2 (-m diamond and –d euk) (51), SignalP 3 (111), and EffectorP v1.01 (64, 112).

### Biological material and molecular biology methods for *Pst* gene expression analysis

We investigated *Pst* gene expression in five different developmental stages or tissue types. We extracted total RNA from dormant spores, germinated spores after 16 hours, 6 and 9 days post infection (dpi) of wheat and from haustoria isolated from wheat leaves at 9 dpi.

In the case of dormant spores, spores were harvested from infected wheat at 14-18 dpi, dried under vacuum for 1 hour and stored at -80°C until use. For germination, fresh spores were heat-treated for 5 minutes at 42^o^C and sprinkled on top of sterile Milli-Q (MQ) water. The container was covered with Clingfilm and spores were incubated at 100% humidity at 10°C in the dark for 16 hours before harvesting. For infection assays, dormant spores were heat treated for 5 minutes at 42°C, mixed with talcum powder (1:7 w/w) and sprayed homogenously with a manual air pump onto seven-day old wheat seedlings wetted with water using a spray bottle. Plants were maintained in a container at 100% humidity in the dark at 10°C for 24 hours. At this point plants were transferred to a constant temperature growth cabinet at 17°C with a 16:8 light cycle. We collected infected wheat leaf samples 6 and 9 dpi. Haustoria were purified from wheat leaves at 9 dpi (113). Infected wheat leaves (~20 g) were surface sterilised with 70% ethanol, washed and blended in 250 ml of 1x isolation buffer (0.2 M sucrose, 20 mM MOPS pH 7.2, 1x IB). The homogenate was passed consecutively through 100 μm and 20 μm meshes to remove cell debris. The filtrate was centrifuged at 1080 g for 15 min at 4°C and the resulting pellets resuspended in 80 mL 1x IB containing 30% Percol (v/v). The suspension was centrifuged at 25,000 *g* for 30 min at 4°C. The upper 10 mL of each tube was recovered, diluted 10 times with 1x IB and centrifuged at 1080 *g* for 15 min at 4^o^C. The pellets were resuspended in 20 mL of 1x IB containing 25% Percoll (v/v) and centrifuged at 25,000 *g* for 30 min at 4°C. The upper 10 mL of each tube was recovered, diluted 10 times in 1x IB and centrifuged at 1080 *g* for 15 min at 4°C. Pellets were stained with ConA-488 to visualize haustoria under the fluorescence microscope. The final pellets were frozen in liquid nitrogen and stored at -80°C prior to RNA isolation.

RNA for all samples was isolated as follows. Total RNA was isolated using the QIAGEN Plant RNeasy kit following the manufacturer’s instructions. Initial RNA quality and purity checks were performed on a NanoDrop ND-1000 UV-Vis Spectrophotometer. Samples were treated with DNase I (New England Biolabs) following the manufacturer’s instructions. Samples were purified using the QIAGEN Plant RNeasy kit following the cleanup protocol, and RNA was eluted from columns in 50 μl of RNase-free water. The concentration and integrity of all final RNA samples were verified on the Agilent 2100 bioanalyzer, using the RNA 6000 nano and pico kits. Three biological replicates were processed.

RNA samples were sequenced at the Ramaciotti Centre (Sydney, Australia) on an Illumina HiSeq 2000 instrument as 100-bp paired end reads. Approximately 10 μg of total RNA per biological sample was processed with the TruSeq RNA Sample Preparation Kit v2.

### Differential expression analysis

We trimmed Illumina RNAseq reads using Trimmomatic v0.35 (92) (ILLUMINACLIP:adapter.fa:2:30:10 LEADING:3 TRAILING:3 SLIDINGWINDOW:4:25 MINLEN:35) and assessed read quality with FastQC v0.11.4 (93). We mapped reads using gene models as a guide using STAR v020201 (114). We first generated a genome reference in the genomeGenerate mode using our gff for gene models (--runMode genomeGenerate –sjdbGTFfile --sjdbGTFtagExonParentTranscript Parent). We mapped our RNAseq reads against this reference using STAR in the alignReads mode (--runMode alignReads --readFilesCommand gunzip –c --outFilterType BySJout --outFilterMultimapNmax 20 --alignSJoverhangMin 8 --alignSJDBoverhangMin 1 --outFilterMismatchNmax 999 -- alignIntronMin 20 --alignIntronMax 10000 --alignMatesGapMax 1000000 --outSAMtype BAM SortedByCoordinate --outSAMstrandField intronMotif --outFilterIntronMotifs RemoveNoncanonical --quantMode GeneCounts). We used featureCounts v1.5.3 and our gene annotation to quantify the overlaps of mapped reads with each gene model (-t exon –g Parent) (115). We identified differentially expressed genes in either haustoria or infected leaves relative to germinated spores (|log fold change| > 1.5 and an adjusted p-value < 0.1) using the DESeq2 R package (116). *k*-means clustering was performed on average rlog transformed values for each gene and condition. The optimal number of clusters was defined using the elbow plot method and circular heatmaps drawn using Circos (117). Scripts regarding the gene expression analysis can be found in the gene_expression folder of the github repo.

We compared the expression pattern of alleles in different clusters (Table S6 and S7) in jupyter notebook Pst_104E_v12_secretome_expression_cluster_analysis_ submission_21092017 in the github repo.

### BUSCO analysis

We used BUSCO2 v2.0 4 beta to identify core conserved genes and to assess genome completeness (32). In all cases we ran BUSCO2 in the protein mode using the basidiomycota reference database downloaded 01/09/2016 (-l basidiomycota_odb9 -m protein). We combined BUSCO identification on primary contigs and haplotigs non-redundantly to asses completeness of the combined assembly. For detail see jupyter notebook Pst_104E_v12_BUSCO_summary_submission_21092017 in the github repo.

### Inter-haplotype variation analysis

We mapped trimmed reads against primary contigs using BWA-MEM v0.7.15-r1142-dirty using the standard parameters (94). We called SNPs with FreeBayes default parameters (118) and filtered the output with vcffilter v1.0.0-rc1 (-f “DP >10” -f “QUAL > 20”) (119). SNP calls were summarized with real time genomic vcfstats v3.8.4 (120).

We aligned all haplotigs to their corresponding primary contigs using nucmer of the mummer package (-maxmatch -l 100 -c 500) (35). We fed these alignments into Assemblytics to estimate the inter-haplom variation for each primary contig – haplotig pairing (34). For this analysis, we used a unique anchor length of 8kb based on the length of identified TEs in our *Pst* assembly and a maximum feature length of 10kb. For consistency, we used nucmer alignments filtered by Assemblytics for the allele status analysis (see below). Analysis and summary of variations is shown in jupyter notebook Pst_104E_v12_assemblytics_analysis_ submission_21092017 and Pst_104E_v12_nucmer_and_assemblytics submission_21092017 in the github repo.

### Allele status analysis

We used proteinortho v5.16 in the synteny mode with default parameters (-synteny) to identify alleles between the primary assembly and haplotigs (53). We parsed the results and defined three major allele status categories as follows. Allele pairs were parsed from the ‘poff-graph’ output file. Inter-haploid genome paralogues were parsed from the ‘proteinortho’ output file and checked for absence in the ‘poff-graph’ output file. Potential singletons were defined as gene models being absent from both of these two output files. Alleles were further subdivided into alleles for which the primary and associated haplotig gene models were located on contigs that aligned with each other at the position of the primary gene model (Figure S2A), alleles for which the primary and associated haplotig gene models were located on contigs that did not align with each other at the position of the primary gene model (Figure S2B), and alleles for which the allele of a primary gene model was not located on a haplotig associated with the respective primary contig (Figure S2C). Potential singletons were screened for being located in regions of the primary assembly that were unphased based on Illumina coverage analysis (see above). Genes located in these regions were defined as unphased and removed from the initial list. All other gene models constitute haplotype specific singletons. Analysis details can be found in the jupyter notebooks Pst_104E_v12_defining_alleles_ submission_21092017 and Pst_104E_v12_missing_allele_QC_ submission_21092017.

### Allele variation analysis

We assessed the variation of allele pairs using three approaches. We calculated the Levenshtein distance (80) on the CDS alignments of two alleles, on the codon based protein alignments and we calculated the dN/dS ratios using these two alignments sets with yn00 paml version 4.9 (81). The CDS of two alleles were aligned using muscle v3.8.31 (121) and codon based alignments were generated using PAL2NAL v14 (122). The Levenshtein distance was calculated in python using the distance module v 0.1.3 (123). Analysis details can be found in the jupyter notebook Pst_104E_v12_post_allele_analysis_ submission_21092017.

### Genome architecture analysis

We used bedtools v2.25.0 (96) and the python module pybedtools (97) to perform various genome analysis tasks. This included the calculation of nearest neighbours using the closest function. Details of the analysis can be found in the jupyter notebooks Pst_104E_v12_post_allele_analysis submission_21092017 and Pst_104E_v12_effectors submission_21092017.

### Orthology analysis of candidate effector analysis

We performed orthology analysis with proteinortho v5.16 (-singles) (53) of all non-redundant candidate effectors with publicly available *Pst* genomes. *Pst*-130 (4) and *Pst*-78 (29) protein sets were downloaded from MycoCosm (05/09/2017) (82). *Pst*-0821, *Pst*-21, *Pst*-43 and *Pst*-887 were downloaded from yellowrust.com (30/03/2017) (5, 124). We performed a similar analysis searching for candidate effector orthologs in *Pucciniales* excluding *Pst* genomes. *Puccinia triticina* 1-1 BBBD Race 1 (29), *Puccinia graminis* f. sp. *tritici* v2.0 (6), *Puccinia coronata-avenae* 12SD80 and 12NC29 (125), and *Melampsora lini* CH5 (126) genomes were downloaded from MycoCosm (05/09/2017). The *Puccinia sorghi* genome (127) (ASM126337v1) was downloaded from NCBI (05/09/2017).

### Data and statistical analysis

We used the python programming language (128) in the jupyter notebook environment for data analysis (129). In particular, we used pandas (130), numpy (131), matplotlib (132) and seaborn (133) for data processing and plotting. Statistical analysis was performed using the Scipy (131) and statsmodel toolkits.

## Acknowledgment

We thank the following colleagues for technical advice; Ying Zhang, Sylvain Forêt, Marcin Adamski, Adam Taranto, and Megan McDonald. We thank Ashlea Grewar for technical assistance with rust multiplication. We thank the following colleagues for feedback on the manuscript; Adam Taranto, Megan McDonald, Sajid Ali, Annemarie Fejer Justesen, and Sambasivam Periyannan. We would like to thank Teresa Neeman from the statistical consulting unit at ANU. We acknowledge support by the Genome Discovery Unit (GDU) providing computing facilities. We thank Ashlea Grewar for technical assistance with rust multiplication.

The work conducted by the U.S. Department of Energy Joint Genome Institute, a DOE Office of Science User Facility, is supported by the Office of Science of the U.S. Department of Energy under Contract No. DE-AC02-05CH11231.

This research/project was undertaken with the assistance of resources and services from the National Computational Infrastructure (NCI), which is supported by the Australian Government.

## Authors’ contributions

BS amplified rust spores, extracted high molecular weight DNA, performed assembly using FALCON-Unzip, performed manual curation of assembly, annotated the genome and proteome, performed all additional bioinformatic analysis except for differential expression analysis of RNA-seq data, conceived study, and wrote the manuscript. JS performed differential expression analysis, contributed ideas to data analysis and wrote manuscript. WC performed pathotyping of the rust strain, amplified rust spores and commented on the manuscript. DG performed infection assays, RNAseq assay, extracted haustoria for RNAseq assay, and commented on the manuscript. MEM contributed ideas to data analysis and commented on the manuscript. JMT and PND contributed ideas to the differential expression analysis and commented on the manuscript. MF contributed ideas to data analysis and wrote the manuscript. RFP provided *Pst* urediniospores and commented on manuscript. JPR contributed ideas to methodological development, data analysis and commented on the manuscript.

## Legends for Supplemental Tables, Figures and Files

**Supplemental Table 1: Summary table of PacBio genome sequencing using 13 SMRT cells**

**Supplemental Table 2: Annotation of gene classes on primary contigs and haplotigs**

For each category the number of proteins and the percentage of proteins with a hit in each category is given. The first number in each column indicates the total number of proteins and the second number the percentage within each category.

**Supplemental Table 3: Summary of inter-haplotype allele analysis**

Summary table describing the allele and conservation state of *Pst*-104E genes. Primary contigs and haplotigs were treated as two representative units for orthology analysis in Proteinortho with synteny flag. The three major categories are highlighted in bold.

In case of alleles they can be subdivided into three categories as illustrated in Supplemental Figure 4. Superscript A, B, C, correspond to Supplemental Figure 4 A, B, C.

The number in brackets for haplotype singletons is the number of singletons given by Proteinortho without filtering for unphased gene models based on genome coverage analysis. True haplotype singletons are located in phased regions of the genome.

**Supplemental Table 4: Number of total proteins and predicted secreted proteins in publicly available cereal rust genomes**

Numbers in brackets indicates the percentage of proteins that are predicted to be secreted in each proteome using SignalP3.

**Supplemental Table 5: Summary table of genes located in indicated expression clusters in regards to EffectorP, nuclear localization and apoplastic localization prediction**

We used EffectorP (64), Localizer (65), and ApoplastP (134) for predictions.

**Supplemental Table 6: Summary of the allele state and orthologous expression patterns of secreted protein coding genes located on primary contigs**

The top half of the table shows how many genes are within each cluster and of those, how many are allelic, inter-haplotype paralogs or singletons.

The bottom half of the table shows how the expression of alleles of genes located on primary contigs cluster in the haplotig gene expression analysis.

**Supplemental Table 7: Summary of the allele state and orthologous expression patterns of secreted protein coding genes located on haplotigs**

The top half of the table shows how many genes are within each cluster and of those how many are allelic, inter-haplotype paralogs or singletons.

The bottom half of the table shows how the expression of alleles of haplotig genes cluster in the primary gene expression analysis.

**Supplemental Table 8: Changes in genome size and contig number at different steps in the assembly process**

**Supplemental Figure 1: Short read genome coverage analysis suggests high levels of haplotype specific sequences**

A) to C) Illumina short read coverage plots for the color-coded genome regions when mapping against primary contigs only. D to G) Illumina short read coverage plots for the color-coded genome regions when mapping against combined primary contigs and haplotigs. The peak at ~120-fold coverage represents diploid genome regions, which are similar for both haplotypes. The peak at ~60-fold coverage represents haploid genome regions, which are specific to one haplotype. Most regions on primary contigs that do not align with a corresponding haplotig display haploid genome coverage (C and G) indicating haplotype-specific sequences. Plots are trimmed to the 99^th^ percentile to remove high coverage outliers such as collapsed repeat regions for visualization purposes.

**Supplemental Figure 2: Over half of the *Pst*-104E genome is covered by repetitive elements**

A) Shows repetitive element annotation on primary contigs (83 Mb) and B) on haplotigs (73 Mb). Top panels show percentage genome coverage for all repetitive elements and different sub-categories. These include transposable elements (TEs) of the Class I (RNA retrotransposons) and Class II (DNA transposons), simple sequence repeats (SSR), and unclassifiable repeats (no Cat). Middle and bottom panels show percent genome coverage of Class I and Class II TEs categorized to class, order, and superfamily levels wherever possible. Repetitive elements were identified using the REPET pipeline and classifications inferred from the closest BLAST hit (see method section).

**Supplemental Figure 3: Approximate age of TEs found on primary contigs**

A) and B) show the estimated age of Class I and Class II elements, respectively, at the superfamily level for TEs with more than 5 copies. C) shows the genome coverage per TE superfamily of elements with an approximate age of less than 100 Mya and more than 50 copies. These TEs are most likely to contribute to current genome evolution. Bars are color-coded according to the order level.

**Supplemental Figure 4: Illustration of the three observed allele configuration categories in the *Pst*-104E haplotype-phased assembly**

A) ‘Classic’ alleles identified by Proteinortho for which the gene models of allele pairs are located in regions of primary contigs and associated haplotigs that overlap in whole genome alignments. B) Alleles detected on non-overlapping associated haplotigs. These are alleles identified by Proteinortho for which the gene models of allele pairs are located in regions of primary contigs and associated haplotigs that do not overlap in whole genome alignments. C) Alleles detected on non-associated haplotigs and primary contigs that might represent long range transpositions. These are alleles identified by Proteinortho for which the gene models of allele pairs are located on haplotigs that are not associated with the primary contig. See also Figure 4 for detailed illustration of one example.

**Supplemental Figure 5: Spatial correlation of AT richness, repeat coverage, candidate effectors and BUSCOs**

Circa plot of the 30 largest primary contigs ordered by size. The total length shown is greater than 50 Mb. See legend for details of each track. Lines with reverse arrowheads illustrate regions in the *Pst*-104E genome where candidate effectors cluster in repeat poor regions. Lines with round arrowheads illustrate regions in the *Pst*-104E genome where candidate effectors appear to cluster with each other close to repeat dense regions. RE is the abbreviation for repetitive element.

**Supplemental Figure 6: Candidate effector genes are clustered**

A) Illustration of the clustering approach. In this example (A) genes 1-3 in the black category are clustered together as each is separated by less than 12 kb to their closest neighbour of the same category. Gene 4 is not part of the cluster as it is more than 12 kb away from gene 3. The clustering algorithm is oblivious to the fact that gene I from the red category is interspersed between genes 2 and 3 of the black category. This cluster thus contain 3 black genes. B) Clustering of equal numbers of genes (n=1444) using a maximum distance cut off of 12 kb. The y-axis represents different gene categories and the x-axis represents the number of genes linked by a minimum distance of 12 kb. The numbers next to the circles indicate the total number of genes in each cluster size bin. The circles are drawn in proportion reflecting this number.

**Supplemental File 1: Summary statistics for all transposable elements identified on primary contigs of the Pst-104E assembly**

Tab separated list per TE as generated by the REPET analysis pipeline see Methods for details.

**Supplemental File 2: Functional annotation of Pst-104E protein coding genes**

Tab separated list of gene IDs followed by annotation term or accession number

**Supplemental File 3: Summary statistics of the allelic variation for each allele pair**

Tab separated list of allele pairs, their codon based Levenshtein distance on the CDS and protein level, the percentage identity on the protein level defined by blast, dN/dS ratio calculated by vn00 and the classification for each primary contig and haplotig protein.

**Supplemental File 4: Allele pairs identified between primary contigs and haplotigs of the *Pst*-104E assembly**

Tab separated list of gene IDs with one pairing per line followed by one of the three classifications: “Classic” alleles, alleles on non-overlapping associated haplotigs, alleles on non-associated haplotigs. See Figures S4 for illustration of these allele classifications.

**Supplemental File 5: Inter-haplotype paralogs of the Pst-104E assembly**

List of gene IDs that we identified as inter-haplotype paralogs.

**Supplemental File 6: Unphased gene loci on primary contigs of the Pst-104E assembly**

List of gene IDs that we identified as being unphased based on primary contigs after coverage analysis of Illumina short read mapping data.

**Supplemental File 7: Singletons of the Pst-104E assembly**

List of gene IDs that we identified as singletons as these lack alleles and inter-haplotype paralogs.

**Supplemental File 8: Single haplotype genes of the Pst-104E assembly**

List of gene IDs that we identified as single haplotype genes. See main text for details.

**Supplemental File 9: EffectorP predicted effector candidates of the *Pst-104E* assembly**

List of gene IDs that we identified as candidate effectors using the machine learning approach EffectorP.

**Supplemental File 10: Expression pattern of the *Pst*-104E secretome**

Tab separated list of gene IDs followed by their expression cluster as shown in Figure 4.

**Supplemental File 11: Non-redundant list of candidate effectors of the *Pst*-104E assembly**

List of gene IDs of all candidate on primary contigs and haplotig effector candidate singletons.

**Supplemental File 12: Parameters of the FALCON assembly pipeline used for the *Pst*-104E assembly**

**Supplemental File 13: Parameters of the FALCON-Unzip processing of the *Pst*-104E assembly**

**Supplemental File 14: Parameters of the REPET Tedenovo repeat prediction of the *Pst*-104E assembly**

**Supplemental File 15: Parameters of the REPET Teanno repeat prediction of the *Pst*-104E assembly**

**Supplemental File 16: Parameters of EvidenceModeler to unify gene predictions of the *Pst*-104E assembly**

**Supplemental File 17: Parameters of PASA pipeline to map *de novo* transcripts on the *Pst*-104E assembly**

